# Ethylenediaminetetraacetic acid promotes the accumulation of nitric oxide

**DOI:** 10.1101/2020.10.05.326413

**Authors:** Shizhong Zhang, Shasha Liu, Shenghui Xiao, Min Li, Jinguang Huang, Kang Yan, Guodong Yang, Chengchao Zheng, Changai Wu

**Affiliations:** State Key Laboratory of Crop Biology, College of Life Sciences, Shandong Agricultural University, Tai’an, Shandong, 271018, P.R China

## Abstract

Ethylenediaminetetraacetic acid (EDTA) is a well-established chelating agent used in industry, agriculture, food, and medicine. However, the analysis of an EDTA-sensitive *Arabidopsis thaliana* mutant revealed that EDTA can significantly promote nitric oxide (NO) accumulation, indicating that EDTA has unexpected biological functions beyond its chelating activity. This finding challenges our current understanding about the effects of EDTA on biological systems.

**One Sentence Summary:** An analysis of ethylenediaminetetraacetic acid (EDTA)-sensitive mutants suggests that EDTA can promote the accumulation of nitric oxide.

## Main Text

Ethylenediaminetetraacetic acid (EDTA) is an organic compound widely used as a chelating agent in industry, agriculture, food, medicine, and environmental protection (*1*). The best known function of EDTA is the ability to form complexes with ions of a variety of metals, including Ca^2+^, Mg^2+^, Fe^2+^, and Zn^2+^ (*2*). Its powerful usage makes us gradually forget the genetic toxicology of EDTA (*3*). For example, exposure of human leucocyte cultures to EDTA (0.1 to 1 mM) leads to increased chromosomal aberrations (*4*), iron(II)-EDTA catalyzes cleavage of RNA and DNA oligonucleotides (*5*), which confirmed the unknown and new biological functions of EDTA. However, the underlying mechanism is largely unclear.

To investigate the biological roles of EDTA, we performed forward genetic screening with an ethyl methanesulfonate-mutagenized *Arabidopsis* M2 population using EDTA-induced inhibition of primary root growth (*6*). One EDTA-sensitive mutant, termed *edta*, was isolated for further analysis. On Murashige and Skoog (MS) medium without EDTA, the root length of *edta* was 75.99% of the wild type (Col-0). With increasing concentrations of EDTA in the medium, the primary root growth of *edta* was significantly inhibited compared to that of the Col-0 plants. For example, at 0.1 mM EDTA, *edta* root length was reduced by 83.78% relative to controls grown without EDTA, whereas Col-0 root length was reduced by only 24.33% (Fig. 1, A and B). *edta* root length was restored on MS medium without Fe-EDTA or EDTA. However, if the medium contained 0.1 mM EDTA, omitting microelements, macroelements, FeSO_4_, CaCl_2_ or MgSO_4_ did not restore the root length of *edta* (Fig.1C, and figs. S1 and S2). In addition, when EDTA was replaced by other chelating agents such as citric acid (CA) (*7*) and sodium dimethyl dithiocarbamate (SDDC) (*8*). *edta* and Col-0 showed similar variation tendency of root length, that is, all showed greatly shortened root length as the concentration of CA and SDD increased (fig. S3). These results demonstrate that the *edta* mutant is highly sensitive to EDTA and imply EDTA has biological functions other than as the chelating agent.

**Fig. 1.**
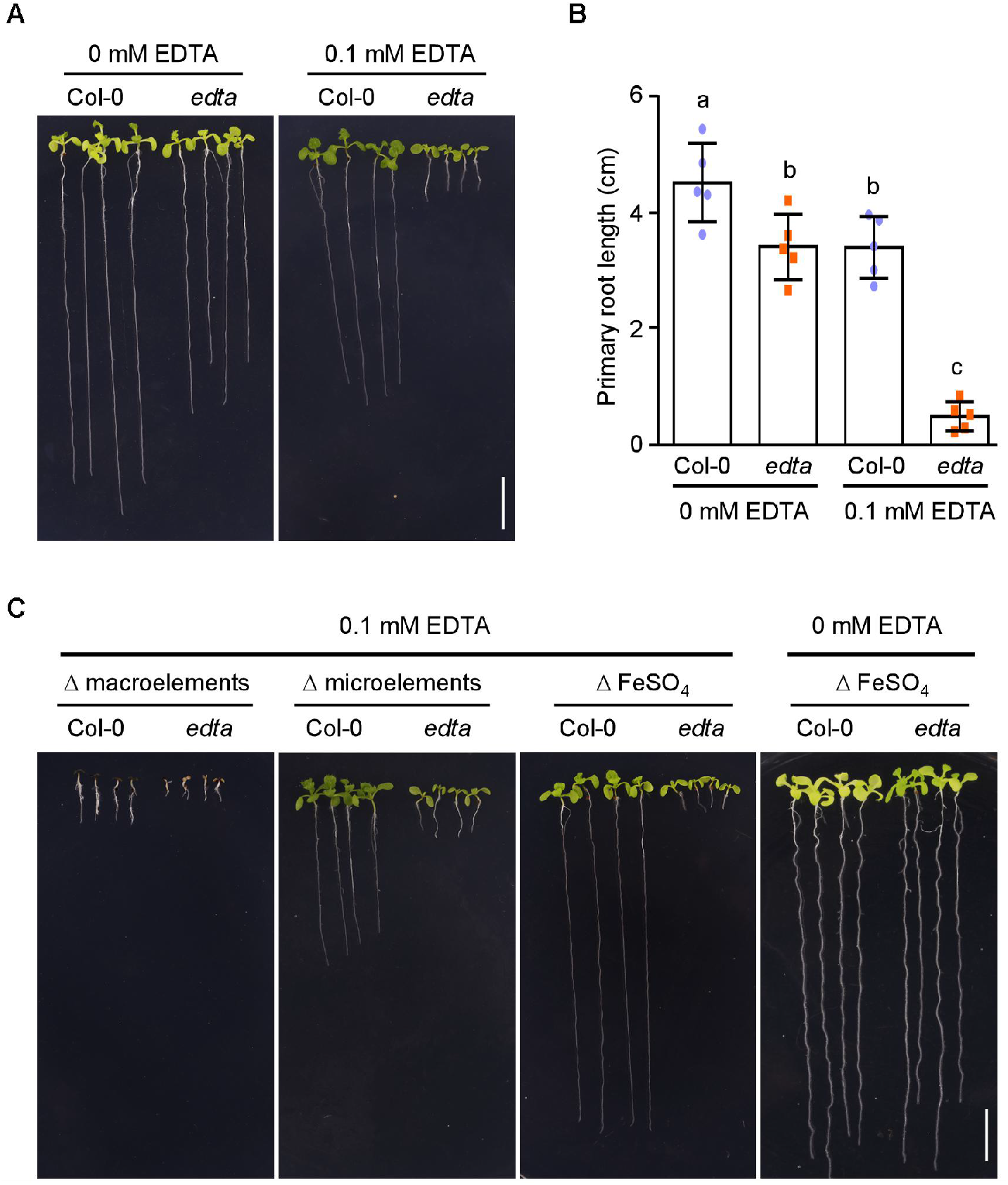
Characterization of *Arabidopsis* EDTA-sensitive mutant. **(A)** The primary root growth of *edta* seedlings is significantly inhibited by EDTA. Col-0 and *edta* seeds were germinated and grown on MS medium containing 0 or 0.1 mM EDTA for 14 d. Scale bar, 1 cm. **(B)** Statistical analysis of primary root length of Col-0 and *edta* in (A). Student’s *t-*test (*P* < 0.05) was used to analyze statistical significance. **(C)** Primary root growth of Col-0 and *edta* seedlings grown on MS medium without microelements, macroelements, FeSO_4_, or Fe-EDTA for 14 d, respectively. “Δ” indicates absence. Scale bar, 1 cm.

To identify the gene that is responsible for the *edta* phenotype, we used map-based cloning with simple sequence length polymorphism (SSLP) markers. We identified a G to A substitution in the sixth exon of *AT5G43940*, which caused an 11-nucleotide deletion in the mature transcripts, leading to a premature stop codon (Fig. 2A). *AT5G43940* encodes a S-nitrosoglutathione reductase (GSNOR). The T-DNA insertion mutant the *gsnor1-3* (GABI_315D11) and the F1 progency of *gsnor1-3* (♂) × *edta* (♀) all showed hypersensitivity to EDTA (Fig. 2, B and C, and figs. S4 and S5). When GSNOR1 (*pGSNOR1::GSNOR1*) was re-introduced into *edta*, the EDTA-sensitive phenotype of *edta* was completely restored. The complementary line (*pGSNOR1::GSNOR1*/*edta*) showed similar EDTA sensitivity as that of Col-0 (Fig. 2, B and C, and figs. S4 and S5). We also assayed the GSNOR activity and established that *edta, gsnor1-3*, and *gsnor1-3edta* had obviously reduced activity compared to Col-0 and *pGSNOR1::GSNOR1*/*edta* (Fig. 2D). Based on these results, we conclude that the EDTA sensitive phenotype of the *edta* mutant was caused by a mutation in the *GSNOR1* gene.

**Fig. 2.**
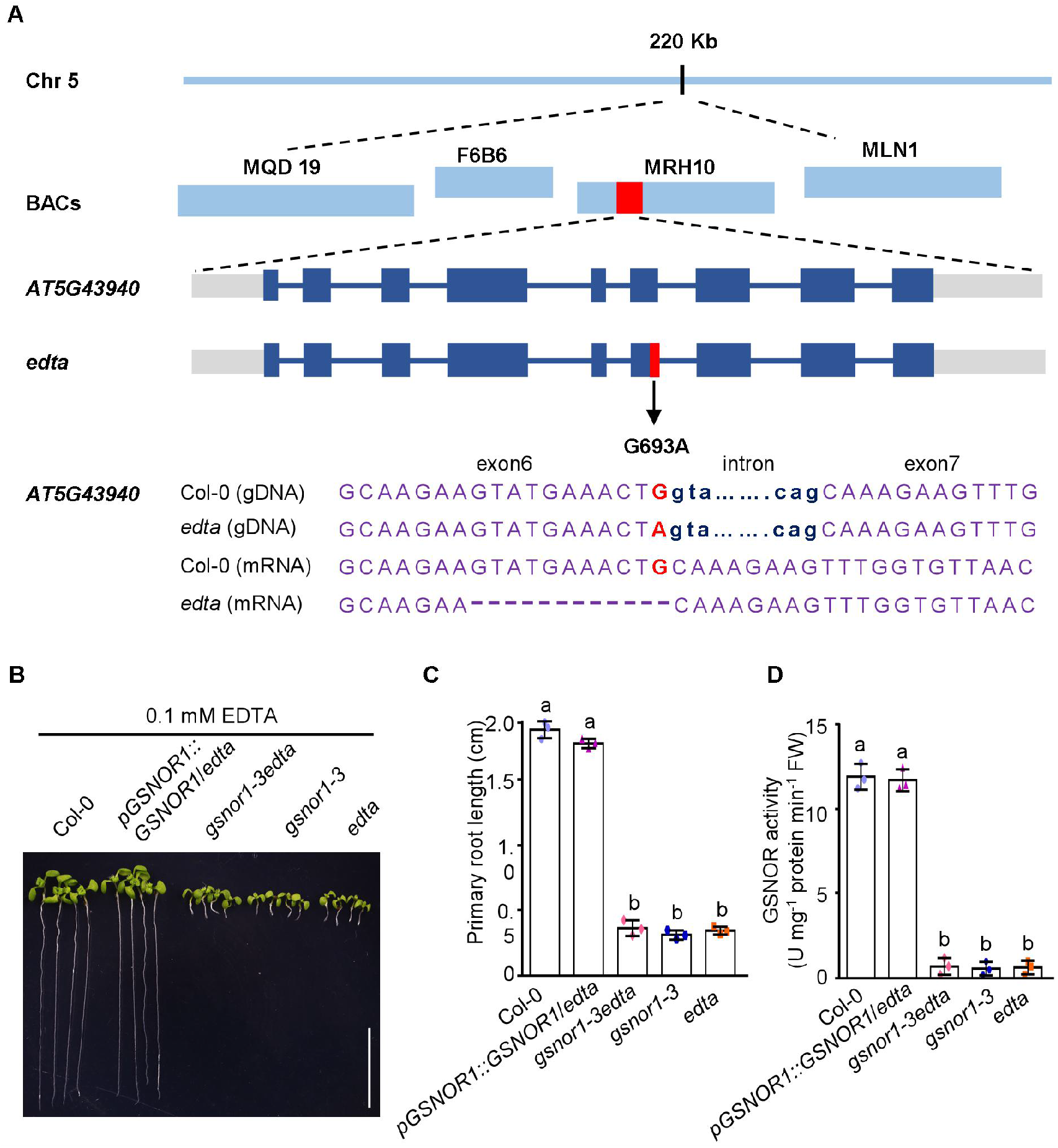
The *edta* phenotype was caused by mutation in *GSNOR1*. **(A)** Map-based cloning of *GSNOR1*. The *GSNOR1* locus was mapped to chromosome 5 between MDQ19 and MLN1, using a total of 2,400 samples. The G693 to A was identified in *AT5G43940* in the sixth exon. Blue line represents chromosome 5, blue bars represent marker on chromosome 5, red bar represents the locus of *AT5G43940* on chromosome 5, gray bars represent untranslational region (UTR), dark blue bars represent exon, red font represents mutant site, purple fonts represent nucleotide sequences. **(B)** Primary root growth of Col-0, *pGSNOR1:GSNOR1/edta, gsnor1-3edta, gsnor1-3*, and *edta* seedlings. *gsnor1-3* is an *GSNOR* null mutant, *gsnor1-3edta* was obtained by crossing *gsnor1-3* and *edta*, and *pGSNOR1:GSNOR1*/*edta* was obtained by expressing the *GSNOR1* cDNA sequence under the control of the *GSNOR1* native promoter in *edta* mutant seedlings, which were germinated and grown on 0.1 mM EDTA (MS) for 10 d. Scale bar, 1 cm. **(C)** Statistical analysis of primary root length in (B). Error bars indicate SEM (*N*=3). Student’s *t*-test (*P* < 0.001) was used to analyze statistical significance. **(D)** GSNOR enzyme activities in Col-0, *pGSNOR1:GSNOR1/edta, gsnor1-3edta, gsnor1-3*, and *edta* seedlings (7 d). Error bars indicate SEM (*N*=3). Student’s *t*-test (*P* < 0.001) was used to analyze statistical significance.

Previous studies indicated that loss function of *GSNOR1* results in higher levels of GSNO (S-nitrosoglutathione, a natural NO donor) and excessive accumulation of NO in plants (*9*). We measured the NO content in roots of *edta, gsnor1-3*, and Col-0 seedlings grown on MS medium with 0, 0.05, 0.1, or 0.2 mM EDTA using the NO-sensitive fluorescent dye 4-amino-5-methylamino-2’,7’-difluorescein diacetate (DAF-FM DA) (*10*). As EDTA concentration increased, NO accumulated in all seedlings, and much higher NO levels were observed in *edta* and *gsnor1-3* than in the Col-0 (Fig. 3, A and B). However, the expression and enzyme activity of *GSNOR1* were not induced by EDTA (fig. S6). So, EDTA can promote NO accumulation and thereby plays an unexpected biological role in NO independent of *GSNOR1*.

**Fig. 3.**
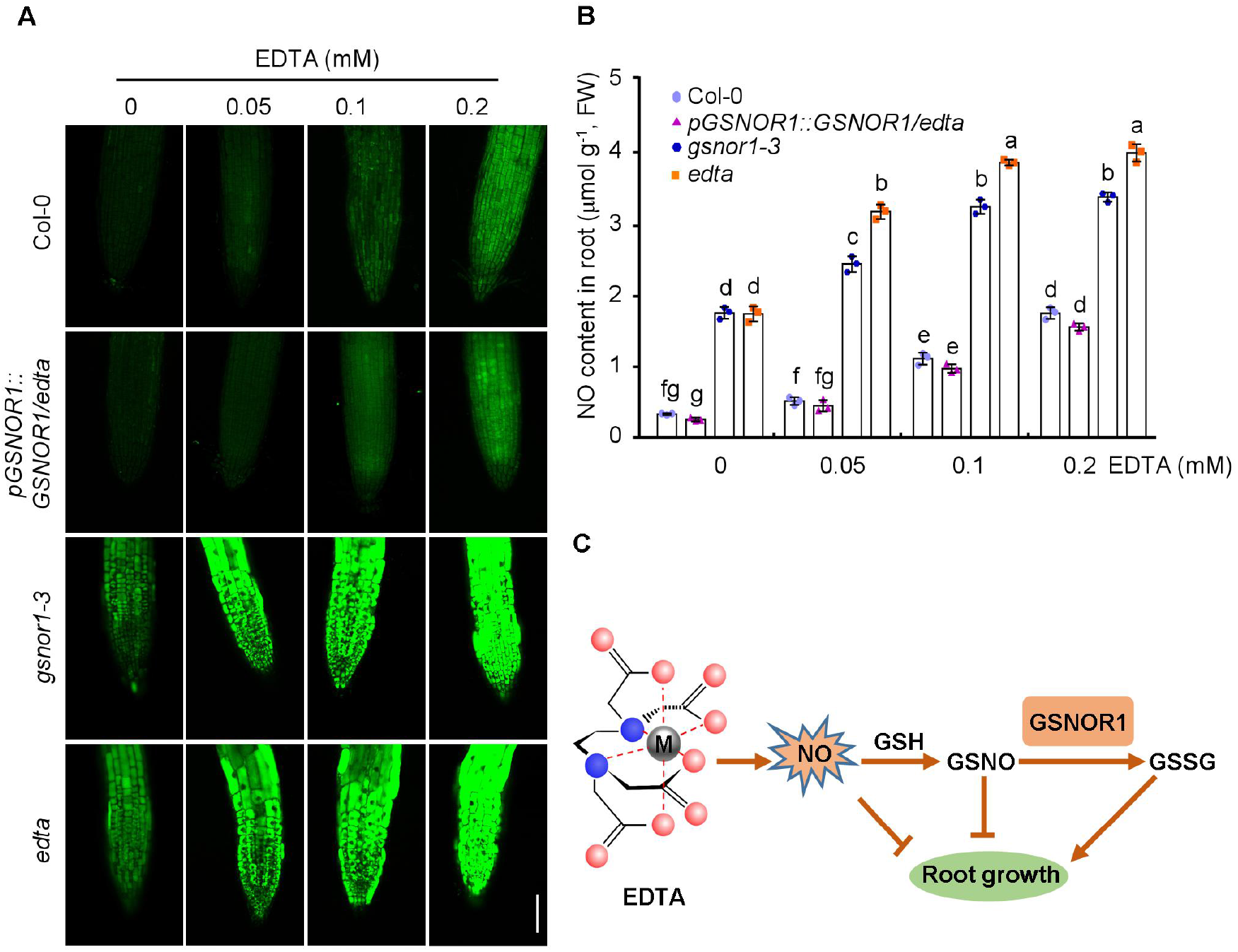
EDTA triggers NO accumulation in *Arabidopsis* seedlings. **(A)** DAF fluorescence in roots of Col-0, *pGSNOR1:GSNOR1/edta, gsnor1-3*, and *edta* seedling grown on 0, 0.05, 0.1, and 0.2 mM EDTA (MS) medium for 7 d. Scale bar, 50 µm. **(B)** Quantitative analyses of NO content in the roots of Col-0, *pGSNOR1:GSNOR1/edta, gsnor1-3*, and *edta* seedlings grown on 0, 0.05, 0.1, and 0.2 mM EDTA (MS) medium for 7 d. Error bars indicate SEM (*N*=3). FW, fresh weight. Student’s *t-*test (*P* < 0.05) was used to analyze statistical significance. **(C)** A simplified biological function of EDTA. EDTA can promote NO accumulation, which can destroy cell structure. Moreover, NO can increase the GSNO and inhibit root growth. The red ball represents oxygen atom, the blue ball represents nitrogen atom, “M” represents metal ion.

NO is an important signaling molecule that regulates growth, development, and stress responses in plants (*11, 12*). In all living organisms, GSNO derived from the reaction of NO and glutathione (GSH), is a major reservoir of bioactive NO species (*13, 14*), in which a NO molecule is covalently attached to a reactive cysteine thiol of a target protein (*15, 16*). GSNO is irreversibly degraded by GSNOR, a class of highly conserved NADH-dependent reductases presented in all living organisms, catalyze the reduction of GSNO to glutathione disulfide (GSSG) and ammonia (NH_3_) in the presence of GSH (*17, 18*). As EDTA can affect specific biological processes via NO, it is possible that EDTA also participates in the protein S-nitrosylation pathway. For example, EDTA might promote S-nitrosylation of PINs (*19*), VND7 (*20*), MC9 (*21*), AHP1 (*22*), and NADPH oxidase (*23*) and thereby negatively regulate auxin signaling, polar auxin transportation, xylem development, cytokinin signaling, cell death in plant immunity, and other processes. In this study, EDTA led to the accumulation of NO, especially in GSNOR inactivated mutants. GSNOR is unable to reduce S-nitrosoglutathione (GSNO), leading to increased nitrosylation of protein and root growth inhibition (Fig. 3C).

Previous studies have reported that EDTA has toxic effects such as chromosomal aberrations, DNA damage, and programmed cell death (*3*). In addition, EDTA could activate the cGMP/PKG/ATP-sensitive potassium channel signaling pathway in mice (*24*) and the MAP kinase ERK in gut epithelial cancer cells (*25*). Therefore, we speculate that these EDTA-activated phenomena may be caused by changes in NO signaling or the protein S-nitrosylation pathway.

These newly uncovered biological effects of EDTA might provide novel interpretation to some previous conclusions. For example, based on a study of the mutant *hot5-2* (*gsnor1-3*), *GSNOR* was identified as a major quantitative trait locus contributing to natural variation of tolerance to Fe toxicity (*26*). However, because iron was supplied as an EDTA chelate, increase of iron concentration meant a concomitant increase in EDTA (*26*). Therefore, based on these data alone, it would not be possible to determine whether *hot5-2* was sensitive to Fe (iron) or EDTA. In current study, we established that *gsnor1-3* displayed hypersensitivity to EDTA or Fe-EDTA rather than to iron. When FeSO_4_ or Fe-EDTA was added to the EDTA-deficient medium, we found that the growth of *edta, gsnor1-3* were inhibited on the medium with FeSO_4_ added alone, and the trend was similar to WT. However, as the concentration of Fe-EDTA increased in the medium, *edta* and *gsnor1-3* did not grow at all (fig.S7). These results indicated that EDTA has more important role in this progress. Another previous study determined that excess iron reduces root tip growth through NO-mediated disruption of potassium homeostasis in *Arabidopsis* (*27*). However, these experiments were also conducted using MS medium (containing EDTA), meaning that excess iron was also present in the excess EDTA treatments (*27*). Given the effect of EDTA on NO accumulation observed in the current study, these results may need to be reinterpreted.

In summary, EDTA exhibits previously unknown biological functions related to NO, which may prompt us to review previous findings, including those concerning the mechanism of metal ion toxicity (*26*). Simultaneously, we should pay special attention to the applications of EDTA in medicine and food, as this chemical may produce negative effects on humans. Even though EDTA is the most widely used synthetic organic compound, the specific mechanism underlying the NO accumulation caused by EDTA remains elusive. In addition, EDTA may exert its biological effects independent of NO signaling, the stories about the underlying mechanism of EDTA have just begun.

## Acknowledgments

We thank Dr. Jianzhong Liu (College of Chemistry and Life Sciences, Zhejiang Normal University, Jinhua, Zhejiang 321004, China) for his kind provision of *gsnor1-3* mutant.

## Author contributions

S.Z., C.W. and C.Z. conceived and designed the experiments; S.Z., S.L., S.X. and M.L. performed the experiments; C.W., S.Z. and S.L. analyzed the data, made the figures and wrote the article; J.H., K.Y., and G.Y. provided important suggestions; C.Z. supervised and complemented the writing. All authors read and approved the final manuscript.

## Data and materials availability

All data are available in the main text or the supplementary materials.

## Supplementary Materials

Materials and Methods

Figures S1-S6

Supplementary Table 1

References (28-31)

