## Supplemental data for "Ethylenediaminetetraacetic acid promotes the accumulation of nitric oxide"

**This PDF file includes:**

Materials and Methods

Figs. S1-S6

Supplementary Table 1

References (28-31)

**Materials and Methods**

EDTA-sensitive mutant screening

Approximately 72,000 EMS-mutagenized *Arabidopsis thaliana* (Col-0) M2 plants were screened for EDTA-sensitive mutants. EMS-mutated seeds were sown on Murashige and Skoog (MS) medium containing 3% (w/v) sucrose, 0.8% (w/v) agar and incubated at 4℃ in the dark for three days. After 7 d of growth, seedlings with severely short roots were transferred to soil. After harvesting, the seeds of these plants were sown on MS medium without EDTA. Plants with recovered root growth were selected as putative EDTA-sensitive mutants.

Plant growth conditions

*Arabidopsis* Col-0 (WT), *edta*, *gsnor1-3*, *gsnor1-3edta*, and *pGSNOR1::GSNOR1/edta* plants were grown under 16 h light at 22℃and 8 h darkness at 22℃. For the root length assays, seeds were sown and grown on differently modified MS medium (pH 5.80-5.85) as indicated, respectively. The 7 to 14-d-old seedlings were photographed and the root lengths were measured using ImageJ. Three biological replicates were performed. *gsnor1-3* is a *GSNOR* null mutant*, gsnor1-3edta* was obtained by crossing *gsnor1-3* and *edta*, *pGSNOR1:GSNOR1/edta* was obtained by expressing the *GSNOR* cDNA sequence under the control of the *GSNOR1* native promoter in the *edta* mutant.

Map-based cloning and mutant complementation

The *edta* mutant was crossed with Landsberg erecta to obtain the F2 population for selection of mutant plants based on increased EDTA sensitivity when grown on MS medium containing 0.1 mM EDTA. Approximately 2,400 *edta* seedlings were selected to map the mutant gene using simple sequence length polymorphism (SSLP) markers (*28*). The mutant gene was initially mapped to chromosome 5 between MDQ19 and MLN1, and the locus was then narrowed down to F6B6. All genes in this region were sequenced, and one mutation, G693 to A693 was identified in the sixth exon of *AT5G43940*, which caused a premature stop codon.

RNA extraction and RT-PCR

Total RNA from 14-d-old seedlings grown on MS medium with 0, 0.05, 0.1, or 0.2 mM EDTA was extracted using TRIzol (Invitrogen, Carlsbad, California, USA) following to the manufacturer's instructions. Reverse transcriptions were performed using Prime Script RT Enzyme MIX I (TaKaRa, Ohtsu, Japan) with oligo dT primer. The cDNA used for RT-PCR was synthesized using the PrimeScriptTM First-Strand cDNA Synthesis Kit (TaKaRa, Ohtsu, Japan). RT-PCR performed as described by C. T. Chen *et al* (*29*). *EF-1α* was used as the internal control. Primers are listed in supplementary Table S1.

GSNOR enzyme activity detection

GSNOR activity was determined as described by B. Gong *et al* (*30*). Approximately 0.1 g of 14-d-old seedlings was collected. The samples were extracted with 0.15 M Tris-HCl (pH 8.0), 25% (v/v) glycerol, and 2% (w/v) PVP40 buffer, then centrifuged at 10,000 g for 20 min at 4℃. The supernatant was incubated at 25℃ with 20 mM Tris (pH 8.0), 200 μM NADH, 400 μM GSNO, and assays were conducted in cuvette with a EVOLUTION 201 UV-Visible spectrophotometer (Thermo Fisher Scientific, USA) by monitoring the formation or decomposition of NADH at 340 nm, after which the decrease in absorbance at 340 nm (dA340/dt) was measured at $min^{-1}$ intervals for 8 mins.

Imaging of NO in *Arabidopsis* roots

The NO in seedling roots was visualized by staining with the NO-sensitive fluorescent dye 4-amino-5-methylamino-2',7'-difluorescein diacetate (DAF-FM DA Molecular Probes, USA). To detect NO, 7-d-old seedlings were loaded with 10 μM of the probe for 30 min at 37℃ in darkness, and washed for 10 min with distilled water. Images were recorded using an LSM880 high-resolution laser confocal microscope (Zeiss, Germany) at 488 nm (*31*).

Measurement of NO content

NO content in the roots was determined using a Micro NO Content Assay Kit (Solarbio, USA). NO content in 7-d-old seedlings (0.2 g per sample) was calculated based on the observation at 550 nm ($OD_{550}$) measured using an ELIASA spectrophotometer (BIO-TEK ELX800, USA) following the manufacturer's instructions.

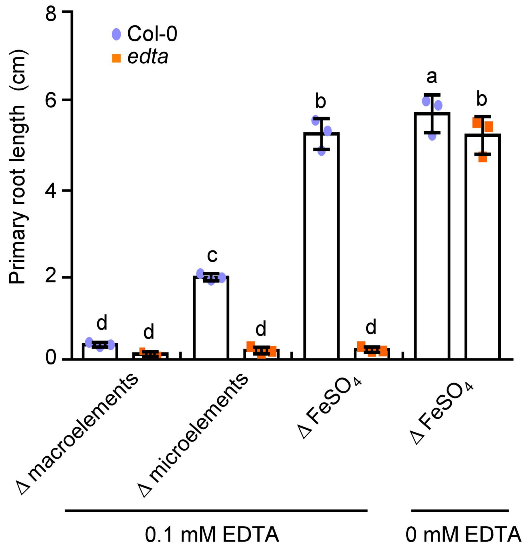


**Fig. S1. Statistical analysis of primary root length of Col-0 and *edta* seedlings grown on MS medium without microelements, macroelements, $FeSO_4$, or Fe-EDTA of Fig. 1C.**

Error bars indicate SEM ($N = 3$). Student's $t$-test ($P < 0.05$) was used to analyze statistical significance.

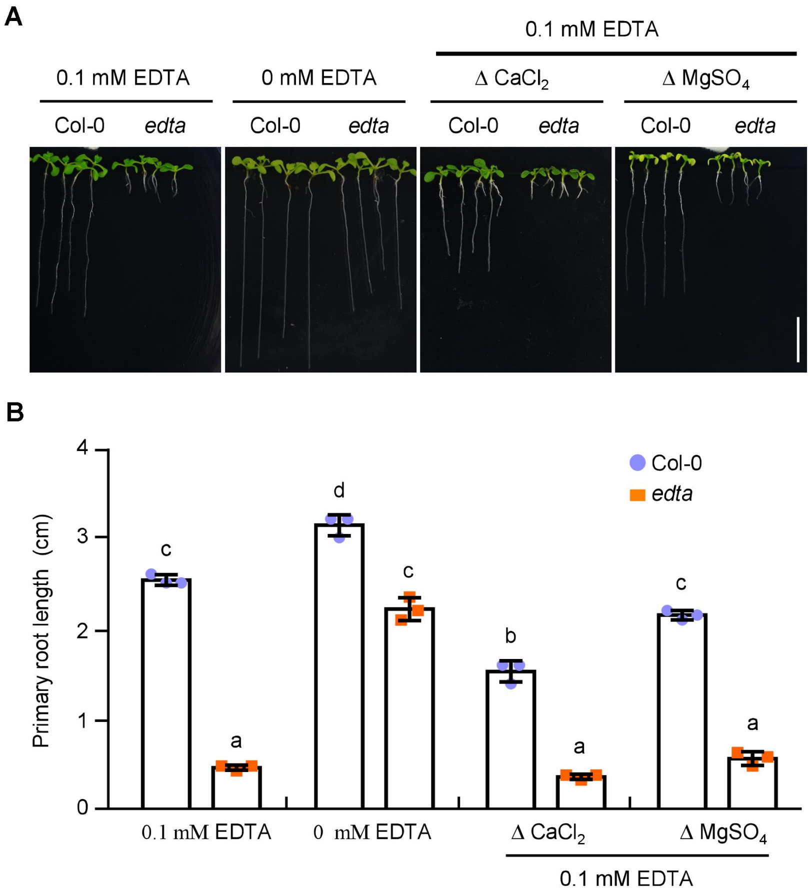


**Fig. S2. The phenotypes of *edta* are not recovered on MS medium without $CaCl_2$ or $MgSO_4$.**

**(A)** Primary root growth of Col-0 and *edta* seedlings grown on MS medium without $CaCl_2$ or $MgSO_4$ for 10 d, respectively. "Δ" indicates absence. Scale bar, 1 cm.

**(B)** Statistical analysis of primary root length of Col-0 and *edta* seedlings in (A). Error bars indicate SEM ($N = 3$). Student's *t*-test ($P < 0.05$) was used to analyze statistical significance.

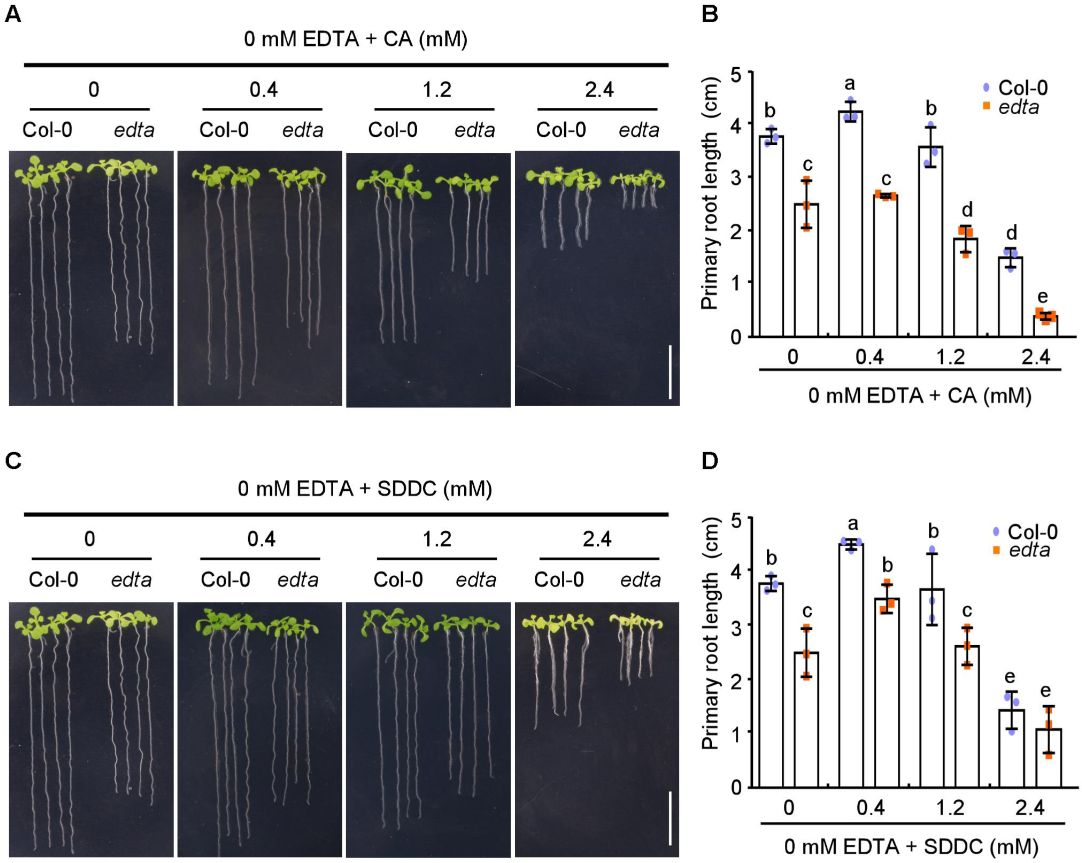


**Fig. S3. *edta* is insensitive to citric acid (CA) and sodium dimethyldithiocarbamate (SDDC).**

(**A**) Primary root growth of Col-0 and *edta* seedlings grown on MS medium supplemented with 0.4, 1.2, or 2.4 mM CA in absence of EDTA for 14 d. Scale bar, 1 cm.

(**B**) Statistical analysis of primary root length of Col-0 and *edta* seedlings shown in (A). Error bars indicate SEM ($N = 3$). Student's $t$-test ($P < 0.001$) was used to analyze statistical significance.

(**C**) Primary root growth of Col-0 and *edta* grown on MS medium supplemented with 0.4, 1.2, or 2.4 mM SDDC in absence of EDTA for 14 d. Scale bar, 1 cm.

(**D**) Statistical analysis of primary root length of Col-0 and *edta* seedlings in (C). Error bars indicate SEM ($N = 3$). Student's $t$-test ($P < 0.001$) was used to analyze statistical significance.

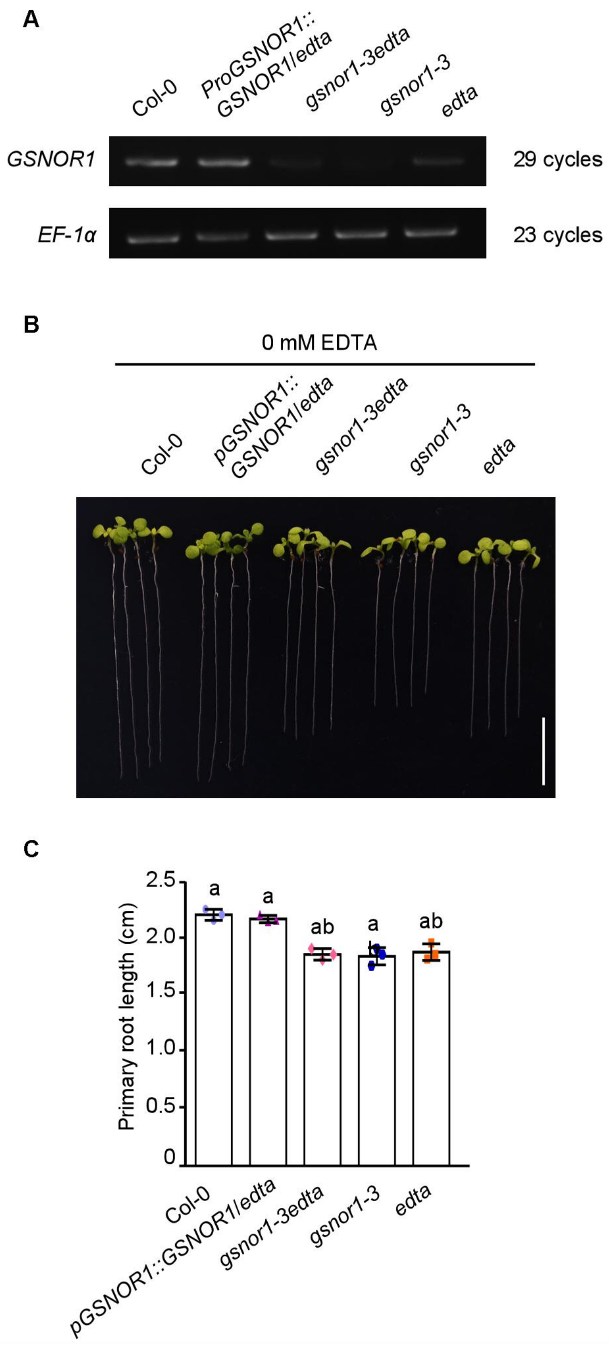


**Fig. S4. The *edta* phenotype without EDTA.**

(**A**) Expression analysis of *GSNOR1* in Col-0, *pGSNOR1:GSNOR1/edta*, *gsnor1-3edta*, *gsnor1-3*, and *edta* seedling by RT-PCR. *EF-1α* was used as a control.

(**B**) Primary root growth of Col-0, *pGSNOR1:GSNOR1/edta*, *gsnor1-3edta*, *gsnor1-3*, and *edta* seedlings grown on MS medium without EDTA for 10 d. Scale bar, 1 cm.

(**C**) Statistical analysis of primary root length of the seedlings in (A). Error bars indicate SEM ($N = 3$). Student's *t*-test ($P < 0.001$) was used to analyze statistical significance.

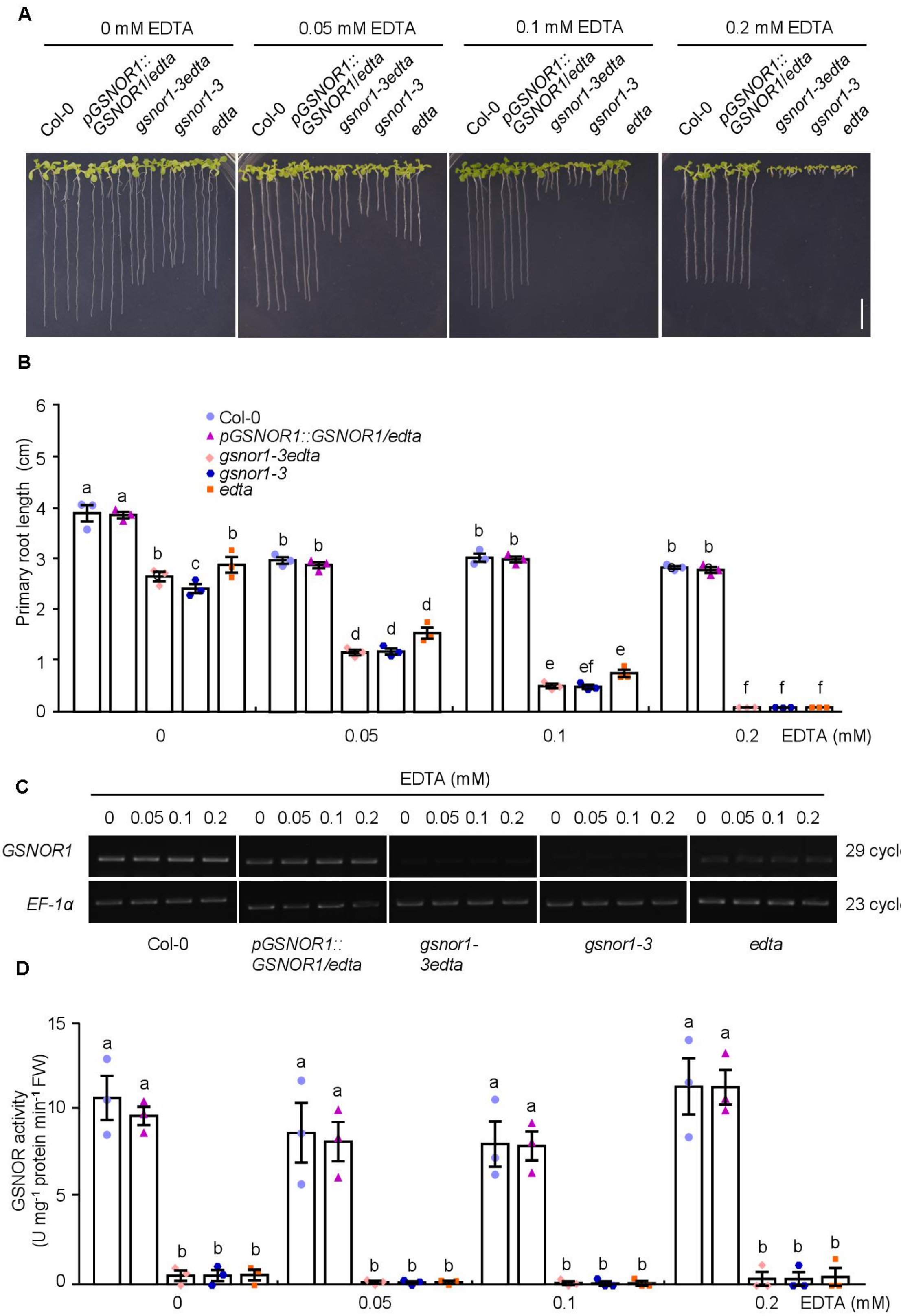


**Fig. S5. The phenotype of Col-0, *pGSNOR1:GSNOR1/edta*, *gsnor1-3edta*, *gsnor1-3*, and *edta* seedlings grown on MS with different concentrations of EDTA.**

**(A)** Primary root length of Col-0, *pGSNOR1:GSNOR1/edta*, *gsnor1-3edta*, *gsnor1-3*, and *edta* seedlings grown on MS containing 0, 0.05, 0.1, or 0.2 mM EDTA for 14 d.

**(B)** Statistical analysis of primary root length of the seedlings in (A). Error bars indicate SEM ($N = 3$). Student's $t$-test ($P < 0.001$) was used to analyze statistical significance.

**(C)** Expression analysis of *AtGSNOR1* in the seedlings in (A) . *EF-1α* was used as a control.

**(D)** GSNOR activities in the seedlings in (A). Error bars indicate SEM ($N = 3$). Student's $t$-test ($P < 0.001$) was used to analyze statistical significance.

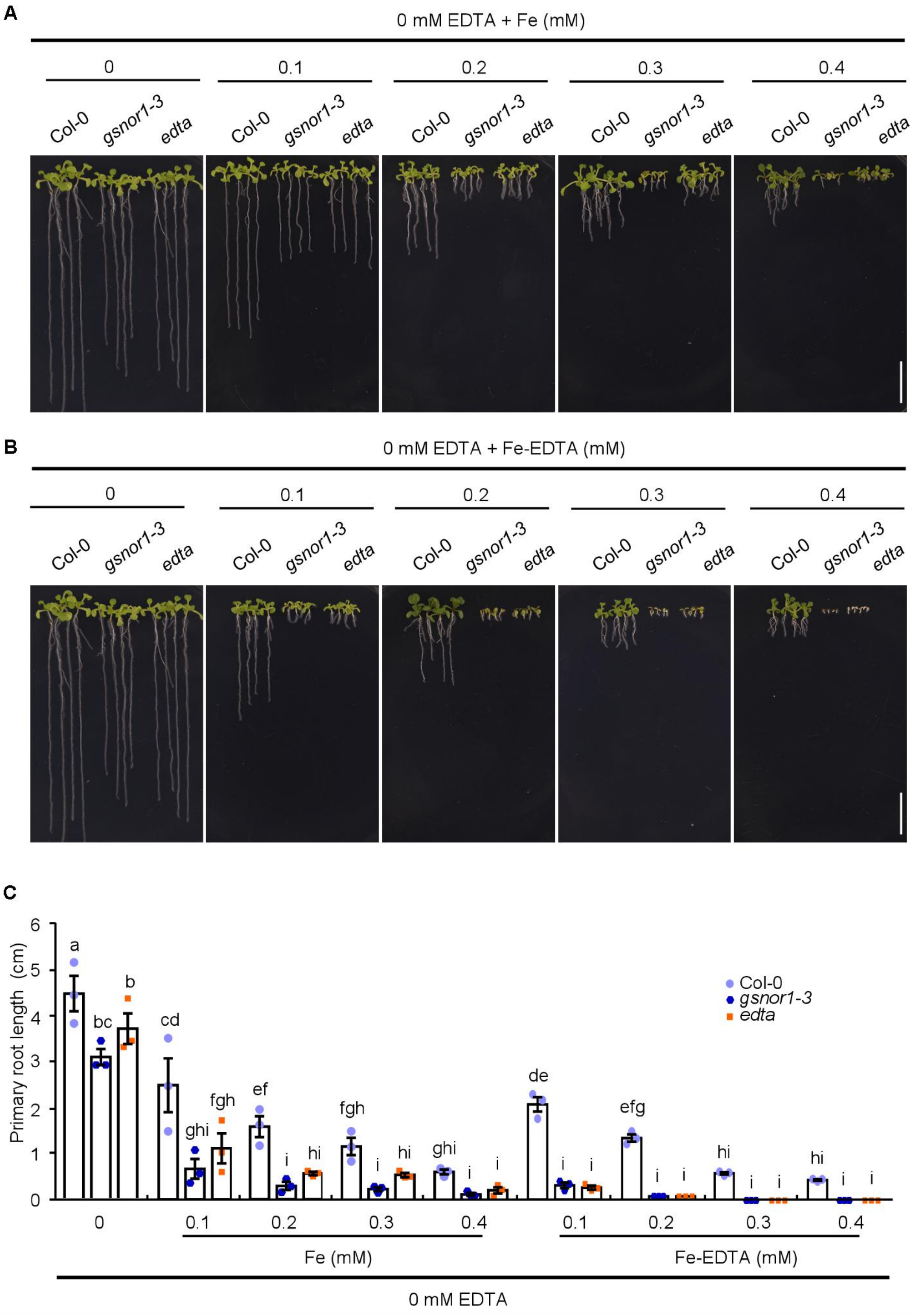


**Fig. S6. The phenotype of Col-0, *gsnor1-3*, and *edta* seedlings grown on MS with different concentrations of $FeSO_4$ or Fe-EDTA.**

**(A)** Primary root growth of Col-0, *gsnor1-3*, and *edta* grown on MS medium supplemented with 0.1, 0.2, 0.3, or 0.4 mM $FeSO_4$ in absence of EDTA for 14 d. Scale bar, 1 cm.

**(B)** Primary root growth of Col-0, *gsnor1-3*, and *edta* seedlings grown on MS medium with 0.1, 0.2, 0.3, or 0.4 mM Fe-EDTA for 14 d. Scale bar, 1 cm.

(**C**) Statistical analysis of primary root length of Col-0, *gsnor1-3*, and *edta* seedlings in (A) and (B). Error bars indicate SEM ($N = 3$). Student's *t*-test ($P < 0.01$) was used to analyze statistical significance.

**Supplementary Table 1. Primers used in this study.**

| Primer | Sequence（5′-3′） | Usage |
|---|---|---|
| 23-19-F | ATAACTTTTTCCTTTTGAAC | Mapping |
| 23-19-R | CATCTTTTTGGGATCTTG | Mapping |
| 23-34-F | ATCACAATGGGTTAAACA | Mapping |
| 23-34-R | CTATATGGCCAGTGAATG | Mapping |
| 23-45-F | CGTGGTCACTTGGATGGT | Mapping |
| 23-45-R | GGCAACAAGACTTCAAGGA | Mapping |
| 23-46-F | CGTTGCTCGTGGATTTTGTAA | Mapping |
| 23-46-R | CTTGTATAAGTTCTTGCCTGTGA | Mapping |
| 23-47-F | TGAATGCTTAGCAAAAAGAA | Mapping |
| 23-47-R | TCGGAAAATTGAGTTTAGTA | Mapping |
| 23-48-F | TGTCCTTCTCTGCGTATG | Mapping |
| 23-48-R | AACCATTGGATATAATTAAG | Mapping |
| 23-51-F | ATTTAGATCGCTTCATTCAC | Mapping |
| 23-51-R | ACCATTTTCACATTTTATTA | Mapping |
| 23-57-F | ACGTTAGTCGTACGGTAGAG | Mapping |
| 23-57-R | TAATTTGCATCGATTTATTA | Mapping |
| 43810-1-F | GGCCAAGGAAGGGATCAGTT | Identification of candidate genes |
| 43810-1-R | CTGTAGCAAGCAAAGAGAGACG | Identification of candidate genes |
| 43810-2-F | CCTAAACAGATCTCTGAATGCTCA | Identification of candidate genes |
| 43810-2-R | GCCGAACTACAAGCAAACAACC | Identification of candidate genes |
| 43890-F | GATTCGATCAAACTCACCCCTAAAG | Identification of candidate genes |
| 43890-R | GGTATGAGCCCTAACTTGAGCC | Identification of candidate genes |
| 43900-1-F | GTCTTCTTGACGTATAAATACGTCTG | Identification of candidate genes |
| 43900-1-R | GCTGCTGACTATTCAGTTTGCC | Identification of candidate genes |
| 43900-2-F | GGATGGTCGAATATTCATTTCGGCTG | Identification of candidate genes |
| 43900-2-R | GTGGCAAGAAGACTAGCAACAGG | Identification of candidate genes |
| 43900-3-F | GACGAAGCCATCCCTTCTTATTATC | Identification of candidate genes |
| 43900-3-R | CAATGAGCATTACCACGGTCAG | Identification of candidate genes |
| 43900-4-F | GTTCAACTCGTCAGAGAACACC | Identification of candidate genes |
| 43900-4-R | GCAGAACGGTGAAAATTCTTAGTTTAAG | Identification of candidate genes |
| 43900-5-F | GGCTGTGCCATCTGCTTATCTG | Identification of candidate genes |
| 43900-5-R | CTGAGCTTCTCAACTTTCAGGAAC | Identification of candidate genes |
| 43940-F | CTCTCTCATTTCTTCCTGCGTC | Identification of candidate genes |
| 43940-R | CATAGATTAAAGCAGAGAGGACCC | Identification of candidate genes |
| 43980-F | CGGAAAACTTTCTTTTTTCCGGAC | Identification of candidate genes |
| 43980-R | TTGGGAAAATAAGAATTGGTAGGT | Identification of candidate genes |
| 43830-F | TTCCCGAGAAAATCATCACTGAG | Identification of candidate genes |
| 43830-R | CGCGTTTAGAACATTGTGGTTTG | Identification of candidate genes |
| 43840-F | GCATCCTCAAGCTTCAGCTATTC | Identification of candidate genes |
| 43840-R | CAATGGAGTGGAAGGTGTGGC | Identification of candidate genes |
| 43850-F | CTGATCTTCATCACTCTGATGAGTCTG | Identification of candidate genes |
| 43850-R | CAACTATCCAGAAGGAGCCGCTG | Identification of candidate genes |
| 43860-F | CACCTCATTCTCATGAACCCACC | Identification of candidate genes |
| 43860-R | CGGGTTAATATCATAAGCAAAGAGGTG | Identification of candidate genes |
| 43870-1-F | CGAAAGCTTCACATAACTTCCCCAC | Identification of candidate genes |
| 43870-1-R | GCGGCGGTCAAAGTCATGATATC | Identification of candidate genes |

| | | |
|---|---|---|
| 43870-2-F | GCGGTTAACGTTCGTTCTGCC | Identification of candidate genes |
| 43870-2-R | GCCCCATTAGTTTCCATATTCATGTG | Identification of candidate genes |
| 43880-1-F | CTTCTGATTTTGTGATGTTTCAGGC | Identification of candidate genes |
| 43880-1-3′ | CTTGTCGATGATTCAGGGAGAG | Identification of candidate genes |
| 43880-2-F | CCTTAGGTGATATGCTTGCACTTCC | Identification of candidate genes |
| 43880-2-R | CACTACTTTCCACTATCAGTCACCC | Identification of candidate genes |
| 43910-1-F | CGAATTTAGCACATAGCCGGAG | Identification of candidate genes |
| 43910-1-R | CCAATTGATCCGTGAATTACAGAAC | Identification of candidate genes |
| 43910-2-F | GTTCTACTAACTTCCCTCAGGTGAG | Identification of candidate genes |
| 43910-2-R | GACTTTGAGATTTTATACGTCAAGAAGAC | Identification of candidate genes |
| 43920-F | CCTTTGTGGACCTTTCAACTATTG | Identification of candidate genes |
| 43920-R | GGAACTACTAGAACTAGGTCCTGG | Identification of candidate genes |
| 5g43940-F | TCTAGACTCTCTCATTTCTTCCTGCGTC | Construction of complementary strains |
| 5g43940-R | GAGCTCCATAGATTAAAGCAGAGAGGACCC | Construction of complementary strains |
| Pr43940-F | AAGCTTGAGTGGTGATGGTTTCAGGGAG | Construction of complementary strains |
| Pr43940-R | TCTAGATGACGCAGGAAGAAATGAGAGAG | Construction of complementary strains |
| S43940-F | GCCTTGGAGCAGTTTGGAATACTGC | Identification of mutant sequence |
| S43940-R | CTTGTGACAGCACTCCAATGCAGC | Identification of mutant sequence |
| 43940-RT-F | CCCTTGTTATCAAGCTGAGTGTCGTGAATGC | RT-PCR |
| 43940-RT-R | CGATGTCAATGCCAATGATCCTTGAAGCACCAGC | RT-PCR |
| EF-1α-F | F:5′-CATCATTTGGCACCCTTCTTCACTGC-3′ | RT-PCR |
| EF-1α-R | R:5′-GTATGGTTGTTACCTTTGCTCCCACAG-3′ | RT-PCR |
